# MetaFunPrimer: primer design for targeting genes observed in metagenomes

**DOI:** 10.1101/2020.07.01.183509

**Authors:** Jia Liu, Paul Villanueva, Jinlyung Choi, Santosh Gunturu, Yang Ouyang, Lisa Tiemann, James R. Cole, Jaejin Lee, Adina Howe

**Author notes:** Address correspondence to Adina Howe. Jia Liu and Paul Villanueva contributed equally to this work. Author order was determined by amount of leading contribution.

## Abstract

High throughput primer design is needed to simultaneously design primers for multiple genes of interest, such as a group of functional genes. We have developed MetaFunPrimer, a bioinformatic pipeline to design primer targets for genes of interests, with a prioritization based on ranking the presence of gene targets in references, such as metagenomes. MetaFunPrimer takes inputs of protein and nucleotide sequences for gene targets of interest accompanied by a set of reference metagenomes or genomes for determining genes of interest. Its output is a set of primers that may be used to amplify genes of interest. To demonstrate the usage and benefits of MetaFunPrimer, a total of 78 HT-qPCR primer pairs were designed to target observed ammonia monooxygenase subunit A (*amo*A) genes of ammonia-oxidizing bacteria (AOB) in 1,550 soil metagenomes. We demonstrate that these primers can significantly improve targeting of *amo*A-AOB genes in soil metagenomes compared to previously published primers.

**IMPORTANCE:** Amplification-based gene characterization allows for sensitive and specific quantification of functional genes. Often, there is a large diversity of genes represented for a function of interest, and multiple primers may be necessary to target associated genes. Current primer design tools are limited to designing primers for only a few genes of interest. MetaFunPrimer allows for high throughput primer design for functional genes of interest and also allows for ranking gene targets by their presence and abundance in environmental datasets. This tool enables high throughput qPCR approaches for characterizing functional genes.

## INTRODUCTION

Diverse microbes in our surrounding environments are key drivers of nutrient cycling and energy necessary for our lives (1–3). To understand the role of these microbes in environments, we characterize their community composition and structure, their diversity, and their function under various conditions. Efforts for characterizing microbiomes have been aided by the development of molecular techniques in combination with sequencing technologies. Specifically, 16S rRNA gene amplicon sequencing has enabled high throughput characterization of taxa or gene composition to inform community structure (4, 5). These sequencing methods are often limited to characterizing phylogenetic markers within a community and are not optimized for characterization of the functional potential of genes within microbial communities.

To characterize the functional roles of microbes, several approaches have been used. One such method is to isolate and enrich representatives of a function of interest to identify and characterize functional traits and their hosts (6, 7). A challenge to this approach is that cultivating microbes from the environment may not represent those found in the environment (8–11). To complement cultivation of isolates, sequencing-based approaches that do not rely on the ability to grow environmental isolates have been used to characterize functional genes (12–14). Specifically, metagenome sequencing of environmental DNA can be used to characterize diverse functional genes in environmental samples. However, it is often the case that these genes make up only a small fraction of the environmental DNA, which can result in a high cost to obtain this functional information (15). Another method to characterize functional genes has been to target amplicons for PCR-based methods. Like 16S rRNA gene sequencing, these methods amplify a specific target gene. All amplicon-based approaches that target genes of interest rely on the ability of primer sets to amplify these genes of interests. These primer sets and their subsequent amplification reactions are most effective if they are both sensitive and specific to target genes of interest.

Many existing primers have been developed based on sequenced genes or genomes (16– 19). The increasing availability of metagenome sequencing provides new opportunities to expand or redesign primers for target genes for gene targets, especially microbes that may not be cultivated or have genomes available (20). PCR-based characterization of functional gene targets has been recently combined with high-throughput qPCR (HT-qPCR) platforms to assay hundreds of genes in a single run. For example, hundreds of primer sets for high-throughput qPCR arrays have been used to simultaneously characterize antibiotic resistance genes in environmental samples (21, 22).

These technologies now enhance our ability to characterize functional genes in the environment. Specifically, by combining the increased availability of metagenomes and the emergence of HT-qPCR platforms, we can scale PCR-based assays for functional genes of interest. Combining these two resources requires the design of appropriate probes, but is limited in the lack of publicly available software that allows users to design environment-specific primers for specific functional genes. Here, we have developed MetaFunPrimer, a pipeline to perform high throughput primer design to target genes of interest existing in metagenome samples. This tool builds upon existing primer design software for developing PCR or qPCR primers, such as Primer3 (23), which can design primers for specific amplification conditions and product length outputs but are limited to a small number of primers and gene targets. MetaFunPrimer designs primers for targeted functional genes and evaluates and prioritizes these primers against hundreds of environmentally abundant functional genes. Here, we demonstrate the use of MetaFunPrimer for designing novel primers for targeting ammonia oxidizing genes previously observed in agricultural soils. While this study focuses on ammonia monooxygenase subunit A gene of ammonia-oxidizing bacteria (*amo*A-AOB) as a specific target gene of interest, MetaFunPrimer is broadly applicable to diverse genes of interest. An online tutorial of the use of MetaFunPrimer is available at *https://metafunprimer.readthedocs.io/en/latest/Tutorial.html*.

The *amo*A-AOB genes were chosen as a target for functional probe design due to its important role in nitrogen cycling. The *amo*A genes encode ammonia monooxygenase, an enzyme that is the main catalyst in ammonia oxidation. Ammonia oxidation is the first and rate-limiting step of nitrification which converts ammonia to nitrite then nitrate, the chemical form of nitrogen that can potentially result in nitrogen loss from in environmental systems (24, 25). Generally, AOB species belong to either beta or gamma subclasses of the class Proteobacteria, with the majority of AOB associated with genera *Nitrosococcus, Nitrosomonas, Nitrosospira* (26, 27). *Amo*A genes have previously been used as functional markers for analyzing AOB diversity (16, 28, 29), and several primer pairs for conserved regions of *amo*A-AOB genes have been previously used for studying its function (16–19). In this study, we use the example of *amo*A*-*AOB genes to demonstrate the usage of MetaFunPrimer. Specifically, we evaluate the diversity of *amo*A-AOB genes in soil metagenomes, evaluate the sensitivity and specificity of previously published probes to detect these genes, and use MetaFunPrimer to design primers for novel gene targets.

## RESULTS

The steps for MetaFunPrimer primer design of *amo*A-AOB genes include: (1) characterization of reference *amo*A-AOB genes; (2) weighting of target genes based on soil metagenomes; (3) design of primers for selected genes; and (4) computational primer evaluation for alignment to target genes (Fig. 1, Table 1).

**TABLE 1.** Data associated with MetaFunPrimer in the design of amoA-AOB genes.

| Data Associated with MetaFunPrimer | Type | Results for our study |
| --- | --- | --- |
| Curated *amoA*-AOB genes from functional gene database | Input | 1205 nucleotide and amino acid sequences |
| Soil metagenomes | Input | 1550 soil metagenomes |
| Optimal clustering similarity found (Recommended by MetaFunPrimer) | Parameter | 96% |
| Gene clusters included (Recommended by MetaFunPrimer) | Parameter | 10 gene clusters |
| Prioritized genes based on input #1 and #2: Total number of soil abundant genes | Output | 720 genes |
| Non-degenerate primers | Output | 78 primer sets |
| Total number of soil metagenome genes targeted by final primer set | Output | 676 (93.89%) |

**Fig. 1.**
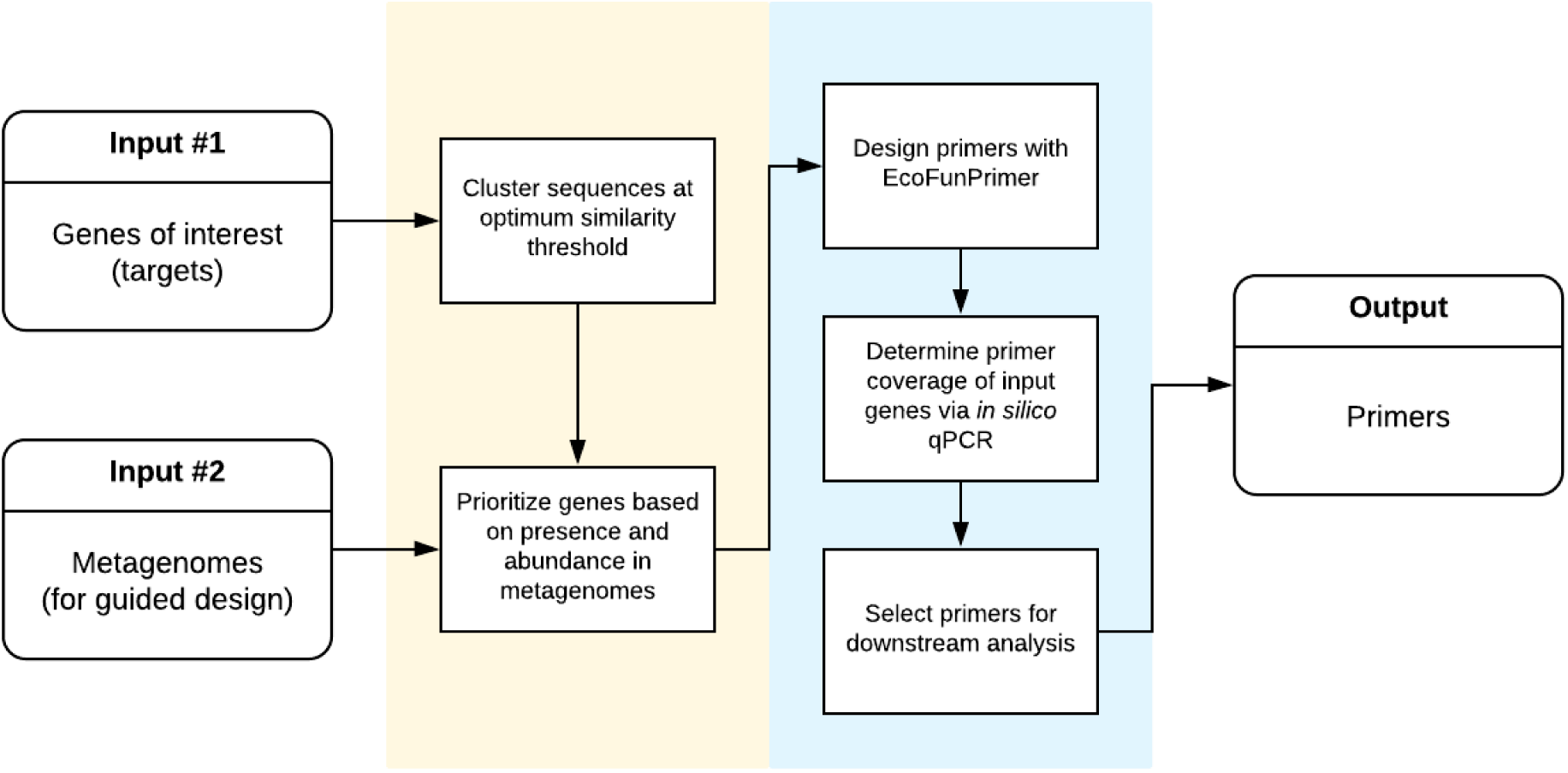
Overview summarizing the MetaFunPrimer pipeline for gene primer design based guided by inputs of reference genes and metagenomes.

### Characterization of reference amoA-AOB genes

A curated set of functional genes for *amo*A-AOB was obtained from the Ribosomal Database Project Fungene (version 9.6) (30). We obtained protein sequences, nucleotide sequences, and their corresponding NCBI accession numbers for a total of 1205 *amo*A-AOB genes. For HT-qPCR applications, we aimed to detect as many target genes as possible with minimal primer pairs. For our study, it was impractical to have thousands of primers, and thus our first step was to reduce the number of gene targets. We removed redundancy and reduced gene targets by initially clustering gene reference sequences based on their similarity. Among the 1205 *amo*A*-*AOB protein sequences, many sequences were observed to have a high degree of similarity. When sequences were clustered from 80 to 100% protein similarity, we found that clustering these sequences at greater than 96% amino acid similarity resulted in the largest increase in resulting total unique clusters (Fig. 2). We aimed to balance the lowest number of clusters representing potential gene targets while representing the most gene diversity. Consequently, we found that clustering based on 96% similarity resulted in a total of 60 clusters, and representative sequences from each cluster covered a wide diversity of *amo*A-AOB including the genera *Nitrosomonas, Nitrosococcus*, and *Nitrosospira* (Fig. S1).

**Fig. 2.**
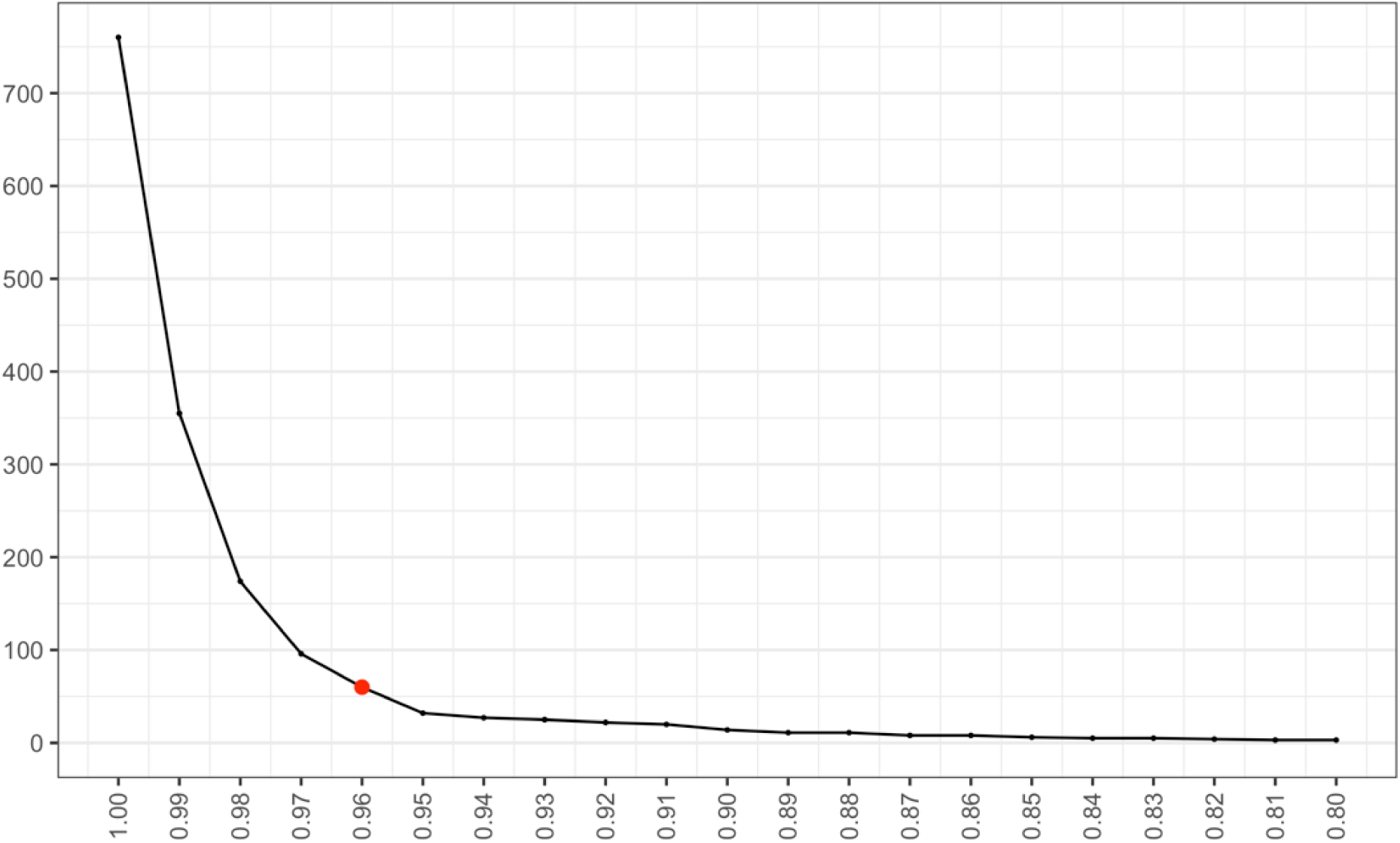
The selection of the appropriate number of genes for designing gene primers can be reduced by clustering sequences by protein similarity. A total of 60 clusters were selected based on 96% amino acid similarity of *amo*A-AOB genes (indicated by red point). Clusters were found using CD-HIT with word size 5 for each of the similarity thresholds indicated.

### Weighting of target genes based on soil metagenomes

The representative protein sequences from each cluster were next aligned to 1550 publicly available soil metagenomes (Table S1), with alignments defined as having 97% percent sequence identity over the length of the reference gene. Each *amo*A*-*AOB associated gene identified in soil metagenomes was then ranked based on two criteria: estimated gene abundance (the total number of observations of each gene within all the metagenomes sequences) and prevalence (the number of unique metagenomes where the gene was observed) (Table S2). The abundance and prevalence of each representative gene were then normalized separately before taking their mean value to calculate each representative sequence’s representation score (R-score). The clusters represented by the ten sequences with the highest R-score accounted for a total of 720 *amo*A-AOB genes, comprising a total of 87.4% of the cumulative overall abundance of these genes observed in the soil metagenomes (Fig. 3).

**Fig. 3.**
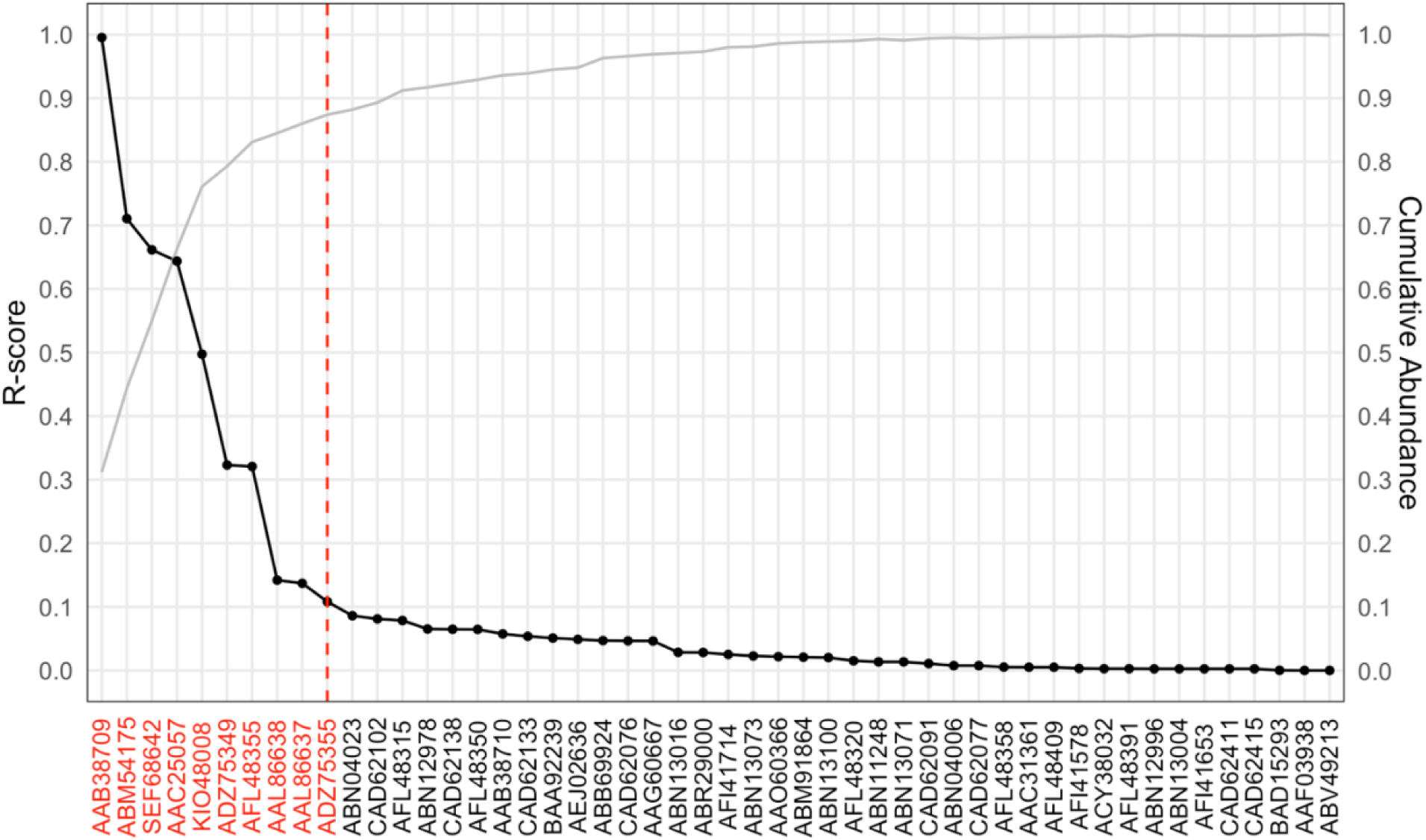
Known *amo*A-AOB genes ranked by representation score (R-score; the mean of the scaled abundance and prevalence) and the estimated cumulative abundance of each gene in 1,550 soil metagenomes. The protein sequences in red indicate those *amo*A-AOB gene clusters and their associated genes that were selected for primer design based on cumulative R-score in reference metagenomes.

### Design of primers for selected genes

The nucleotide sequences of these 720 genes were obtained and used for further primer design. Embedded in MetaFunPrimer is EcoFunPrimer, which was developed by the Ribosomal Database Project (RDP) at Michigan State University (https://github.com/rdpstaff/EcoFunPrimer). EcoFunPrimer is a primer design tool which outputs primers based on input genes. For the 720 genes selected for primer design, 28 primer sets were generated by EcoFunPrimer, allowing at most 6 degenerate primers based on specific PCR conditions (Table S3). From the resulting 28 degenerate primer pairs, MetaFunPrimer generated 181 single non-degenerate primer pairs and next evaluated these primers through an *in silico* PCR against the 720 targeted reference genes. In some cases, redundant primer pairs exist for the same gene target, and these redundant primers were removed resulting in a final set of 78 non-degenerate primer pairs (Table S4). Overall, the resulting primer pairs were predicted to *in silico* amplify a total of 676 out of 720 soil abundant *amo*A-AOB genes observed from soil metagenomes.

Finally, to compare our designed primers to previously published primers, we summarized previously published *amo*A-AOB primers (16–19) to single non-degenerate primer pairs (Table S5). MetaFunPrimer’s *in silico* amplification procedure was performed using these primer pairs to evaluate their alignment to the 720 targeted soil abundant *amo*A-AOB genes. In total, 49.44% (356/720) of these genes would be detected using pre-existing primer pairs, while the primers designed by MetaFunPrimer resulted in 93.89% (676/720) detection (Table 2). Within each soil abundant cluster, primers designed using MetaFunPrimer tend to have higher amplification abilities compared with pre-existing primers.

**TABLE 2.** Comparison of previously published *amo*A-AOB primers to those in this study. Targeting rate is the ratio of the number of genes within the associated cluster that can be aligned by given primer sets and the total number of genes in the cluster.

| Soil abundant *amoA*-AOB cluster [gene representative] | Number of gene sequences within each cluster | Number of previously published primer pairs that target each cluster | Targeting rate of previously published primers | Number of MetaFunPrimer primer pairs that targeting each cluster | Targeting rate of MetaFunPrimer primers |
| --- | --- | --- | --- | --- | --- |
| 1 [AAB38709] | 20 | 3 | 3 (15.00%) | 5 | 19 (95.00%) |
| 3 [SEF68642] | 285 | 7 | 55 (19.30%) | 33 | 273 (95.79%) |
| 4 [KIO48008] | 320 | 14 | 255 (79.69%) | 26 | 304 (95.00%) |
| 5 [AAC25057] | 65 | 11 | 30 (46.15%) | 12 | 53 (81.54%) |
| 6 [AAL86637] | 5 | - | - | 2 | 3 (60.00%) |
| 7 [AAL86638] | 11 | 2 | 10 (90.91%) | 2 | 11 (100.00%) |
| 28 [ABM54175] | 2 | - | - | 2 | 2 (100.00%) |
| 29 [ADZ75349] | 8 | 2 | 3 (37.50%) | 3 | 7 (87.50%) |
| 52 [AFL48355] | 2 | - | - | 2 | 2 (100.00%) |
| 58 [ADZ75355] | 2 | - | - | 1 | 2 (100.00%) |
| Total | 720 | 20 | 356 (49.44%) | 78 | 676 (93.89%) |

Primer pairs that target genes in each cluster are described in Table S5 in supplementary materials.

## DISCUSSION

Amplicon-based approaches for characterizing functional genes provide an approach that is a strong complement to metagenome sequencing. In comparison to metagenome sequencing, HT-qPCR approaches have the potential to be more affordable and sensitive due to the targeted amplification of genes of interest and can be used for standardized surveys of microbial communities and their functions (31). The opportunities of HT-qPCR approaches and amplicon-based approaches depends strongly on the reliability of primer design to target genes of interest (32). In this present work, we introduce the MetaFunPrimer pipeline for designing HT-qPCR primers and demonstrate its use by capturing a broad diversity of relevant genes associated with ammonia oxidation within soil metagenomes. Nitrogen cycling genes are one of the most challenging targets for amplicon approaches as they are encoded by highly diverse microorganisms, including heterotrophic nitrifying microorganisms, denitrifying bacteria, anammox bacteria, nitrifying archaea, and denitrifying fungi (33). Previously, there have numerous efforts to design primers for *amo*A and other nitrogen cycling genes, but existing primers detect a limited range of the phylogenetically diverse genes and often result in misinterpretation (34). Our analysis supports these previous observations that currently existing primers capture less than half *amo*A-AOB genes in soil metagenomes. Using MetaFunPrimer, we have developed 78 novel primer sets to improve quantification of these genes in soil metagenomes, increasing detection of *amo*A-AOB genes from 49% to 94% coverage of observed genes in metagenomes. Notably, in soil metagenomes, *amo*A-AOB genes comprise less than 0.002% of reads in metagenome libraries and thus comprise only a fraction of each generated metagenome. In contrast, qPCR-based approaches would allow for amplification of these genes from environmental DNA, allowing for more sensitive detection.

In our *amo*A-AOB example, we aimed for hundreds of primer sets to capture high diversity of these genes in soils. Generally, however, MetaFunPrimer inputs can be used to design primers for any user-inputted number of sequences, and this number could be varied to suit experimental capabilities or user-specific aims. Another important attribute of MetaFunPrimer is the ability to rank primer design based on targets present in metagenomes. This feature allows for the selection of the most relevant genes based on previous observations of abundance and prevalence in reference metagenomes. For our study, we weighted equally both abundance and prevalence, but the weights of each category could be varied to prioritize diversity or representation within metagenomes. Additionally, the selection of metagenomes as a reference for selecting probes can also be varied. For example, one could use inputs of metagenomes from only bioenergy-associated soils to prioritize microbial communities within specific agricultural sites. Alternately, genomes could be used as a reference for probe design, allowing users to weight primers for genes from known representatives.

Overall, we developed the MetaFunPrimer pipeline as a high-throughput primer design software to partner with the availability of HT-qPCR capabilities. However, this tool is appropriate for any targeted amplification approach, where primer design for specific genes of interests can be guided by available datasets, as we demonstrated in a recent paper which designed primers with the same approach and successfully measured microcystin-producing genes in hundreds of lake water samples (Lee et al., 2020). Within MetaFunPrimer, we also make available workflows for *in silico* comparisons of primers and gene targets. Similar to any primer design effort, experimental validation is required, but computational efforts can help determine which candidates to test experimentally.

## MATERIALS AND METHODS

As inputs, MetaFunPrimer takes the nucleotide and protein sequences of the genes of interest, a file containing the mapping between a gene’s nucleotide and protein sequence, and gene sequences for prioritization (such as metagenomes). The output of the pipeline is a set of primers that can be used to amplify selected functional genes. The major steps of MetaFunPrimer are firstly to filter and rank genes of interest based on both diversity and representation in inputs, and then to design and evaluate primer sequences for genes of interest (Fig. 1).

### Identifying environmentally representative gene clusters and determine target genes

The first step in the MetaFunPrimer pipeline is to cluster input protein sequences over a range of similarity thresholds in order to determine an optimal or user-defined similarity threshold. Specifically, CD-HIT (35, 36) is used to cluster sequences in the range of 80% to 100% (with 1% increments) similarity to determine the number of clusters found at each threshold. MetaFunPrimer will recommend a similarity threshold that optimizes the first-order difference, a criterion based on the symmetric derivative (37). However, users can select the most appropriate cluster similarity threshold based on their needs.

Next, MetaFunPrimer evaluates the presence of these genes in user-input reference sequences, i.e., metagenomes. For each cluster, the representative protein sequence (identified by CD-HIT) is aligned to reference sequences using DIAMOND (version 0.9.14) (38). Each representative protein sequence is then ranked based on their R-score in reference sequences (i.e., in the case of our case study, these are soil metagenomes). The R-score is defined as the mean of that gene’s normalized abundance and prevalence among reference sequences. The representative genes for each cluster of sequences are subsequently ranked based on R-score in descending order, gene clusters are included until the user-input threshold of the cumulative R-score (i.e., 80% in the case study) is reached. Genes that are associated with selected ranked clusters are considered as genes of interests and consequently target genes for primer design and are converted into their corresponding nucleotide sequences.

### Designing and evaluating primers for genes of interest

MetaFunPrimer uses selected gene sequences and user-defined parameters such as amplicon product length and melting temperature ranges for the subsequent primer design process. Within MetaFunPrimer, EcoFunPrimer is the primary tool used to design thermodynamically stable primer pairs from aligned nucleotide sequences. Depending on user-defined inputs, it is possible for primer outputs from this pipeline to have multiple degenerate forms. To evaluate primer effectiveness, MetaFunPrimer converts all primer outputs to non-degenerate forms (e.g., all possible primer pairs) of forward and reverse primers. Next, all primer pairs are evaluated via *in silico* PCR against the original set of reference genes provided by the user. A pair of primers successfully amplifies a gene product if both the forward and reverse primers achieve a 100% match against a sequence. In some cases, a single reference gene may be targeted by multiple pairs of primers, and each primer pair can also potentially target more than one gene. Thus, as a final step, MetaFunPrimer outputs the minimal number of primer sets to achieve a maximum number of reference gene products.

### Data availability

For *amo*A-AOB primer design, 1205 protein and nucleotide sequences and a file containing the mapping between each gene’s nucleotide and protein sequence obtained curated gene sequences from the Fungene database, requiring a Hidden Markov Model (HMM) search score > 400 and HMM coverage over 70.2% amino acid similarity. To prioritize these gene targets for *amo*A-AOB function in soils, we used 1550 publicly available soil metagenomes (Table S1) as reference metagenomes for primer design.

## ACKNOWLEDGEMENTS

This work was funded by the DOE Center for Advanced Bioenergy and Bioproducts Innovation (U.S. Department of Energy, Office of Science, Office of Biological and Environmental Research under Award Number DE-SC0018420). Any opinions, findings, and conclusions or recommendations expressed in this publication are those of the author(s) and do not necessarily reflect the views of the U.S. Department of Energy. This work was conducted under the MMPRNT project, funded by the DOE BER Office of Science award DE-SC0014108.

